# A simple bioreactor-based method to generate kidney organoids from pluripotent stem cells

**DOI:** 10.1101/237644

**Authors:** Aneta Przepiorski, Veronika Sander, Tracy Tran, Jennifer A. Hollywood, Brie Sorrenson, Jen-Hsing Shih, Ernst J. Wolvetang, Andrew P. McMahon, Teresa M. Holm, Alan J. Davidson

## Abstract

Kidney organoids generated from human pluripotent stem cells have the potential to revolutionize how kidney development and injury are studied. Current protocols are technically complex and suffer from poor reproducibility and high reagent costs restricting scalability. To overcome these issues, we have established a simple, inexpensive and robust method to grow kidney organoids in bulk from human induced pluripotent stem cells. Our organoids develop tubular structures by day (d) 8 and show optimal tissue morphology at d14. A comparison with fetal human kidney suggests that d14 organoid renal structures most closely resemble ‘capillary loop’ stage nephrons. We show that deletion of *HNF1B,* a transcription factor linked to congenital kidney defects, interferes with tubulogenesis, validating our experimental system for studying renal developmental biology. Taken together, our protocol provides a fast, efficient and cost-effective method for generating large quantities of human fetal kidney tissue, enabling the study of normal and aberrant human renal development.

## Introduction

The use of human pluripotent stem cell-derived organoids as *in vitro* models to study development and recapitulate disease has been a major advance in the field in recent years (Lancaster and Knoblich, 2014). Organoids provide an advantage over monolayer cell culture due to their more complex multi-cellular 3D architecture that better approximates *in vivo* tissues. With regards to the kidney, traditional monolayer cultures have had limited utility for modeling the structure and function of nephrons (the tubular units responsible for filtering the blood and maintaining salt and fluid homeostasis). This is not surprising, given that nephrons are subdivided into functionally distinct portions that act in series to generate the urine. Specifically, the renal corpuscle, containing interdigitating podocytes wrapped around a capillary tuft, filters the plasma and generates an ultrafiltrate that travels via the proximal tubule, descending and ascending limbs of the Loop of Henle, and more distal segments (distal convoluted tubule, connecting tubule) to the collecting duct (McMahon, 2016).

Early protocols for converting pluripotent stem cells into renal cells attempted to mimic the developmental signals governing early kidney formation from the intermediate mesoderm. A cocktail of secreted factors including low doses (3 μM) of the small molecule Wnt agonist CHIR99021 (CHIR) were found to induce intermediate mesoderm-like cells (Mae et al., 2013). However the major breakthrough in the field was the discovery that higher concentrations of CHIR (8 μM) followed by fibroblast growth factor 9 (FGF9) (Barak et al., 2012, Takasato et al., 2014), a growth factor required for nephron progenitor growth *in vivo,* was sufficient to induce kidney organoids with a global transcriptional profile similar to 1^st^ trimester fetal kidneys (Takasato et al., 2015). Subsequent methodologies showed that FGF9 can be substituted for B27, a serum-free supplement used to maintain neurons in cell culture (Freedman et al., 2015). In addition, more complex combinations of Activin A, Bone morphogenetic proteins (BMPs), and FGF9 have been used to achieve kidney organoid formation (Morizane et al., 2015, Taguchi et al., 2014). How these factors induce a kidney program in pluripotent stem cells remains poorly understood, although it has been suggested that high levels of CHIR may mimic the Wnt signals that induce mesoderm and dominate in the posterior portion of the intermediate mesoderm where nephron progenitors arise (Takasato and Little, 2016). A major drawback of the current protocols is the large cost of the reagents, with FGF9 and B27 being prohibitively expensive, thus severely limiting the large-scale culture of kidney organoids.

Despite commonalties in the factors used to induce kidney organoid formation, the existing protocols differ in their technical details and are overall quite complex. For instance, the method of Taguchi et al. (2014) requires co-culture with spinal cord explants (as a source of Wnt), that of Takasato et al. (2015) involves repeated CHIR treatments, cell dissociation and then re-aggregation on transwell filters, while Freedman et al. (2015) generate epiblast ‘spheroids’ that are cultured in a Matrigel sandwich. How reproducible these protocols are between different pluripotent stem cells lines has been highlighted as a concern (Morizane and Bonventre, 2017a, Cruz et al., 2017) and as patient-derived and gene-edited induced pluripotent stem cell (iPSC) lines from diverse backgrounds continue to be produced, a protocol with universal utility is highly desirable.

In order to overcome the issues with current methodologies, we have developed a novel strategy for generating kidney organoids from iPSCs that is simple, robust, cost-effective and allows large-scale organoid production. Our method involves the formation of embryoid bodies (EBs; balls of embryonic tissue) in the presence of CHIR, followed by culture in medium supplemented with ‘KnockOut Serum Replacement’ (KOSR) as a substitute for FGF9. Importantly, this approach is inexpensive and highly efficient, allowing kidney organoids to be grown in spinner-flasks. The organoids develop nephrons containing podocytes, proximal and distal tubule segments, collecting ducts, endothelial cells and interstitial cells that are comparable to structures reported using other methods. Comparison with human fetal kidneys reveals that our organoid nephrons most closely correspond to the ‘late capillary loop’ stage of differentiation. Prolonged culture of our kidney organoids is associated with an expansion in interstitial cells and their conversion into pro-fibrotic myofibroblasts, suggesting that this may be a new model to study renal fibrotic scar development. We further show that our protocol can robustly generate organoids of equivalent quality from three differently reprogrammed human iPSC lines without a need for line-specific optimization. Finally, we show that disruption of *HNF1B,* a gene implicated in congenital kidney malformations in humans, results in agenesis of kidney tubules, highlighting the value of our kidney organoid method for the study of human kidney development and modeling of renal birth defects.

## Results

### A simple, cost-effective method to generate kidney organoids

We developed a protocol to differentiate human iPSCs into kidney organoids in large numbers via an EB formation step using CHIR, KOSR and simple spinner-flask bioreactors (Figure 1A). Specifically, the initial setup of the assay involves growing iPSC colonies in a monolayer to 40-50% confluency (Figure 1B). On day (d) 0, the colonies are enzymatically detached and mechanically fragmented into pieces by pipetting, and transferred into ultra-low attachment 6-well plates for suspension culture. A 3-day treatment window with 8 μM CHIR in BPEL medium (a fully defined serum-free differentiation medium; Ng et al., 2008) was found to be sufficient to induce aggregation of the colony fragments into spherical EBs (Figure 1C). We discovered that culturing the CHIR-induced EBs in DMEM with 15% KOSR (‘Stage II’ medium) from d3 onwards, supported tubule formation that was visible under brightfield microscopy by d8 (Figure 1D). Gene expression analysis over the course of d1-8 suggests that our approach recapitulates the developmental program of mesoderm formation and nephrogenesis, with downregulation of mesodermal genes by d4 that coincides with a rise in renal progenitor markers (Figure S1). As spinner-flask bioreactors have been found to improve oxygen and nutrient perfusion, and supports organoid differentiation in other contexts (Pagliuca et al., 2014, Martin and Vermette, 2005), we transferred the organoids into 150 ml spinner-flasks at d8. We found that spinner-flask culture of kidney organoids allows concurrent growth and maturation of hundreds of organoids containing tubular kidney tissue by d14 (Figure 1E). This approach is cost-prohibitive using other existing methods, but is feasible in our method due to the low reagent cost of the KOSR/DMEM medium.

**Figure 1.**
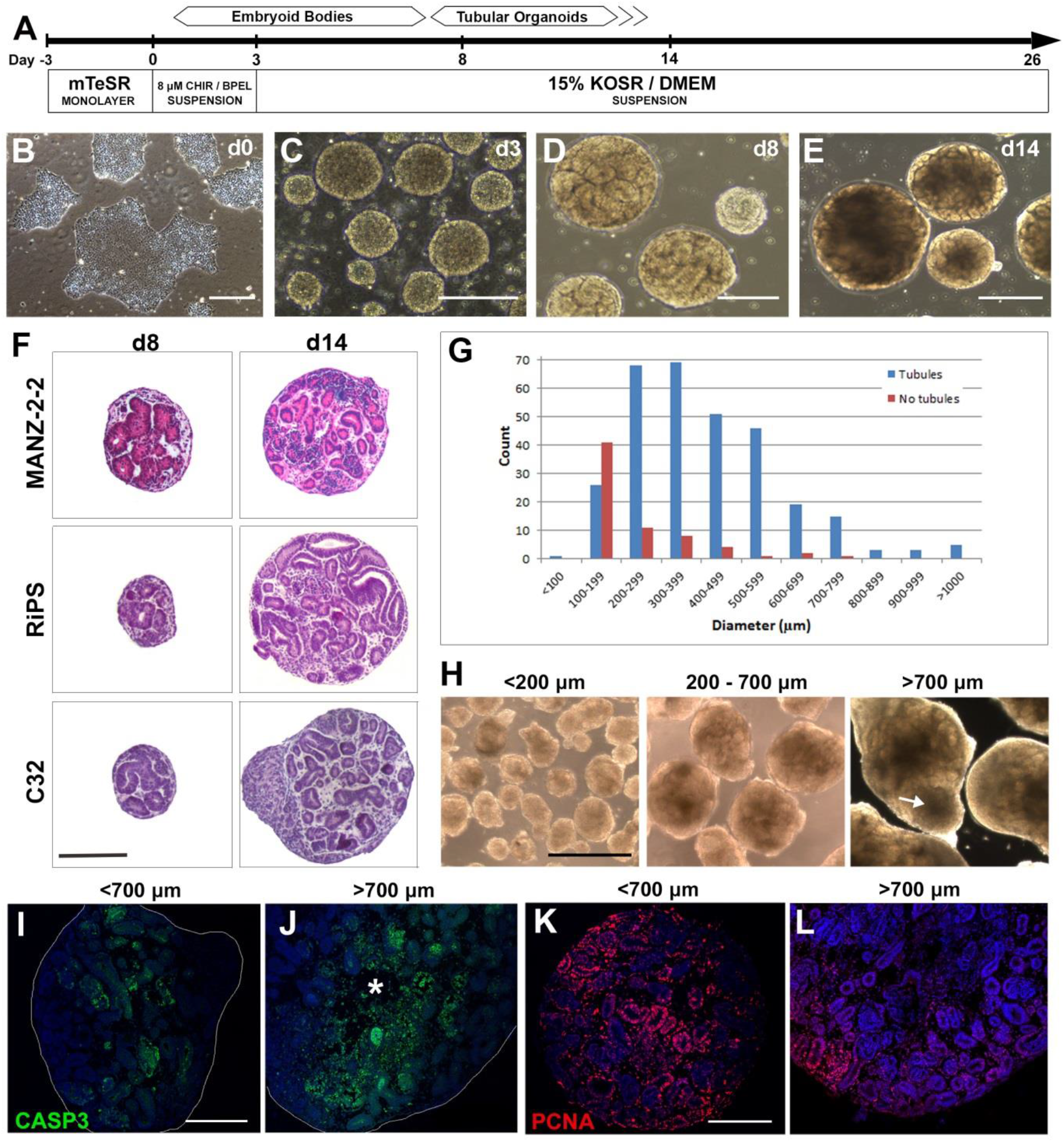
Derivation of kidney organoids from three iPSC lines. **(A)** Overview of the protocol. **(B)** Starting iPSC colonies. **(C)** Embryoid bodies. **(D, E)** Tubular organoids at day (d) 8 and d14. **(F)** Hematoxylin and Eosin stained organoid sections derived from three independent iPSC lines at d8 and d14. **(G)** Bar graph showing the relationship between organoid size, number and presence of tubules in a single representative assay at d14 (n=374 organoids). **(H)** Panels show brightfield images of organoids in the <200, 200-700 and >700 μm size ranges. A non-tubular outgrowth is indicated by the arrow. **(I-L)** Immunofluorescent staining of sections from ‘small’ (<700 μm) and ‘large’ (>700 μm) organoids showing apoptotic cells stained for activated-CASP3 (green) and the proliferation marker PCNA (red). Asterisk in (J) marks a central region with reduced cellularity. Nuclei are stained blue with DAPI. Scale bars: 200 μm (C, D, F, I-L), 400 μm (E), 500 μm (B, H).

Cross-sectioning and histological staining of d8 and d14 kidney organoids confirmed the presence of tubular epithelia, interstitial cells and regions resembling primitive glomeruli (Figures 1F, S2). Comparable results were obtained when the protocol was applied to three independently reprogrammed iPSC lines: MANZ-2-2, RiPS and CRL1502-C32 (herein referred to as C32), although the C32 line had more of a propensity to form non-renal outgrowths (Figure 1F). In terms of efficiency (number of organoids that contain nephron-like structures), we found no major differences between the iPSC lines with kidney organoids comprising 80-90% of the total number of organoids at d8 (n=271/293 for RiPS; n=469/553 for MANZ-2-2, n=848/989 for C32).

We next analyzed whether there was a correlation between organoid size (diameter) and the presence of tubules at d14. We found that ~60% (n=41/68) of the organoids with diameters <200 μm did not form renal structures, while ~90% of the organoids with diameters ≥200 μm contained tubules (n=279/306; Figures 1G, H). This suggests that there is a size threshold that is required before tubules can form. As our protocol involves growing the tissue in suspension we were able to introduce a simple size filtration step using a 200 μm strainer to eliminate these smaller organoids that lacked renal differentiation potential. On the other end of the size spectrum, ~7% (27/374) of the organoids reach a diameter >700 μm (Figure 1H), possibly due to insufficient fragmentation of the iPSC colonies at d0 or due to EBs fusing together. Given that the diffusion of oxygen and nutrients likely becomes limiting as organoid diameter increases (Hubert et al., 2016, Winkle et al., 2012), we examined whether apoptosis was occurring in d14 organoids in the 200-700 μm and >700 μm size ranges. To this end, we performed immunohistochemistry for the apoptosis marker activated caspase 3 (CASP3) on d14 sections of 200-700 μm organoids. While few CASP3^+^ cells were found scattered throughout the tissue and around clusters of presumptive podocytes in 200-700 μm organoids (Figure 1I), >700 μm organoids exhibited a notable loss of cellularity in their cores (asterisk in Figure 1J) and displayed qualitatively more CASP3^+^ cells throughout the tissue. As a further assessment of organoid viability, we performed immunohistochemistry for the proliferation marker, PCNA. While proliferating cells were widespread throughout 200-700 μm organoids (Figure 1K), in >700 μm organoids, cell division was largely restricted to peripheral regions (Figure 1L). Together, these observations indicate that large organoids (>700 μm) are associated with central apoptosis and reduced proliferation, suggesting that they may not be as viable as smaller organoids. Based on this, we incorporated an additional size filtration step using a 500 μm strainer to remove large organoids which, together with the 200 μm strainer, enable collection and culture of a uniform population of optimally viable organoids.

### Kidney organoids contain ‘capillary loop’ stage nephrons

To evaluate nephron differentiation in d14 organoids we next surveyed a range of markers by immunohistochemistry. We identified interstitial cells that stained with a MEIS1/2 antibody and CD31^+^ endothelial cells that were scattered throughout the organoid (Figures 2A, B). The endothelial cells were often found in close proximity to the primitive glomeruli, which contained cells that stained for the podocyte marker Wilms’ Tumor suppressor-1 (WT1) and NPHS1/Nephrin (Figure 2B and data not shown). Electron microscopy revealed that the podocytes display an immature appearance at d14 with broad areas of cell-cell contacts. Prolonged culture of the organoids up to d26 resulted in more elaborate cell-cell junctions resembling the early stages of foot process formation and interdigitation (Figures 3A – B’).

**Figure 2.**
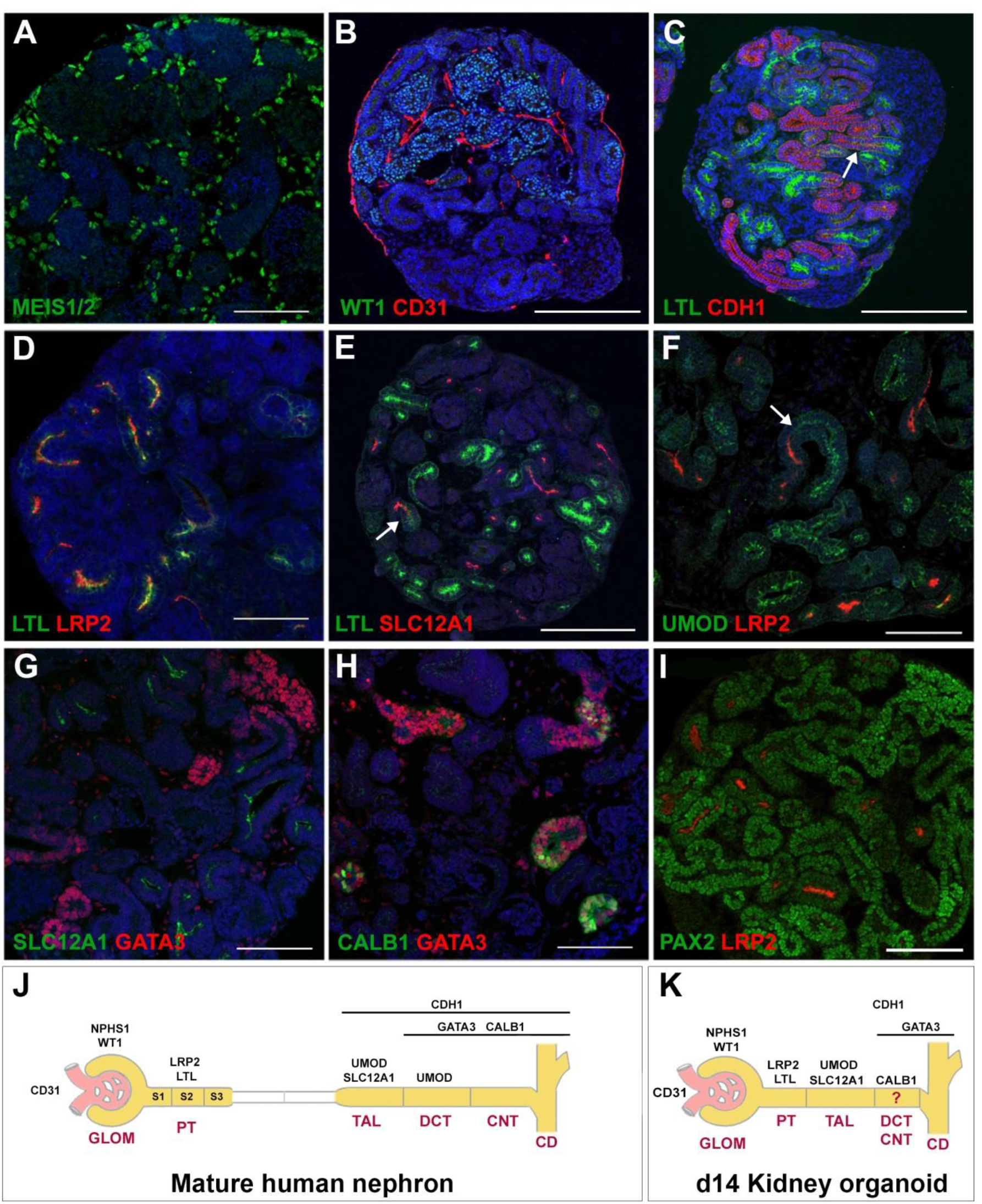
Renal marker analysis in kidney organoids. **(A-I)** Immunofluorescent staining of paraffin sections of d14 organoids showing **(A)** MEIS1/2^+^ interstitial cells (green). **(B)** WT1^+^ podocytes (green) and CD31^+^ endothelial cells (red). **(C)** LTL^+^ proximal tubules (green) and CDH1^+^ distal tubules and collecting ducts (red). **(D)** Co-labeling of LTL (green) and LRP2 (red) in proximal tubules. **(E)** LTL^+^ proximal tubules (green) and SLC12A1^+^ thick ascending limb segments (red). **(F)** LRP^+^ proximal tubules (red) and UMOD^+^ thick ascending limb segments (green). **(G)** SLC12A1^+^ thick ascending limb segments (green) and GATA3^+^ tubules/ducts (red). **(H)** Co-staining of GATA3 (red) and CALB1 (green) in the distal nephron/collecting duct. **(I)** Co-staining of PAX2 and LRP2. **(J, K)** Schematic representations of the segmentation patterns of mature human nephrons (J) and that proposed for d14 kidney organoids (K). Abbreviations: CD, collecting duct; CNT, connecting tubule; DCT, distal convoluted tubule; GLOM, glomerulus; PT, proximal tubule; TAL, thick ascending limb. Nuclei are stained blue with DAPI. Arrows indicate junctions between segments. Scale bars: 100 μm (A, F-I), 200 μm (BE).

**Figure 3.**
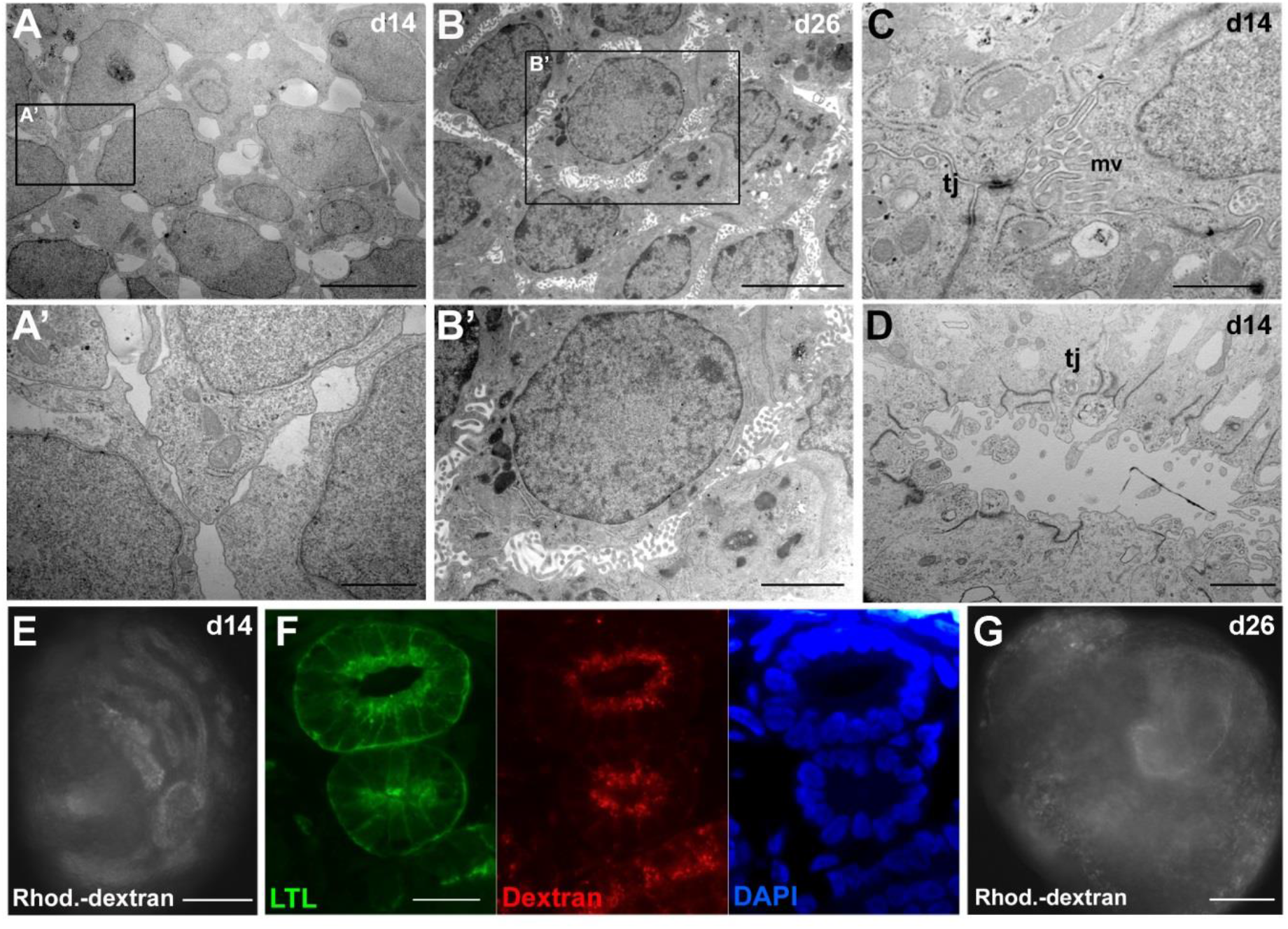
Ultrastructure and tubular absorption of kidney organoid nephrons. (**A-B’**) Transmission electron micrograph (TEM) of presumptive podocytes at d14 (A) and d26 (B; boxed areas are shown in A’ and B’, respectively). (**C**) TEM showing rudimentary microvilli (mv) and tight junctional complexes (tj) in presumptive proximal tubule cells. (**D**) TEM of presumptive collecting duct. (**E**) Fluorescent whole-mount image of a d14 kidney organoid incubated with 10kDa dextran-Rhodamine showing accumulation of dextran in tubules (representative of n=40 organoids). **(F)** Cross-section of a kidney organoid following 10kDa dextran-Rhodamine incubation showing uptake of the dextran (red) into LTL^+^ proximal tubules (green; representative of n=8 organoids). **(G)** Fluorescent whole-mount image of a d26 kidney organoid incubated with 10kDa dextran-Rhodamine showing poor dextran absorption (representative of n=34 organoids). Scale bars: 1 μm (A’, C), 2 μm (B’, D), 5 μm (A, B), 20 μm (F), 100 μm (E, G).

In d14 organoids, developing kidney tubules were found to be subdivided into proximal segments that label with the pan-proximal tubule marker Lotus tetragonolobus lectin (LTL) and more distal portions that label with Cadherin 1/E-Cadherin (CDH1; Figure 2C). In general, there was little overlap of LTL and CDH1 except in short stretches at their junction and only in occasional tubules (arrow in Figure 2C and Video S1 of a serial z-stack). LTL staining was largely restricted to the apical side of the tubular epithelium (in agreement with its localization in kidney tissue sections; Hennigar et al., 1985), but also weakly detected on the basolateral surface of renal epithelial cells. As mature proximal tubules are subdivided into S1-S3 domains, we co-stained the organoids for LTL and LRP2/Megalin, the latter of which is predominantly expressed in the S1 and S2 segments in mature nephrons (Christensen et al., 1995). We found that the staining patterns of LTL and LRP2 fully overlap, indicating that the proximal tubules have not undergone any further sub-segmentation at this stage (Figure 2D). Consistent with this relatively immature differentiation state of the proximal tubule, only a limited degree of apical microvilli were detected by electron microscopy (Figure 3C). Despite these immature features, incubation of the organoids with 10 kDa Rhodamine-labeled dextran for 48 hours results in the specific uptake of the dextran into LTL^+^ tubules, indicating that the absorptive function of the proximal tubule is acquired early in nephrogenesis (Figures 3E, F).

Next, we explored the identity of the distal portion of the nephron. Morphologically, there was no indication of descending or ascending thin limbs, which form part of the loop of Henle and are positioned between the proximal tubule and the thick ascending limb segment. In agreement with this observation, we found that LTL^+^ cells abut a short segment that stains for SLC12A1 and Uromodulin (UMOD), markers expressed by the thick ascending limb (Figures 2E, F; arrows). While we were unable to detect the distal convoluted tubule marker SLC12A3 at this stage, we did find positive staining for GATA3 and Calbindin 1 (CALB1), which label the distal convoluted tubule, connecting tubule, and cortical collecting duct in mature adult nephrons (note: during kidney organogenesis GATA3 is a marker of the collecting duct; Figures 2G, H; Labastie et al., 1995, Kirk et al., 2010, Mantilla et al., 2017). Interestingly, CALB1 was found to label an internal subdomain of the GATA3+ epithelium, possibly reflecting a nascent distal tubule segment (distal convoluted or connecting tubule) connected to a GATA3+ collecting duct (Figure 2H). At the ultrastructural level, the distal nephron/presumptive collecting duct tubules display large lumens made up of columnar epithelial cells with well-defined cell junctional complexes and few apical microvilli (Figure 3D).

To further assess the differentiation state of d14 organoid nephrons we examined PAX2, as this transcription factor is largely downregulated following terminal differentiation of renal epithelial cells (Dressler and Woolf, 1999). We found low levels of PAX2 in the proximal tubules but slightly higher levels in the distal nephron/presumptive collecting duct (Figure 2I). Collectively these data show that d14 kidney organoids, in comparison to mature human nephrons (Figure 2J), are comprised of immature nephrons made up of a primitive glomerulus, an LTL+/LRP2+ proximal tubule segment attached to an SLC12A1+ thick ascending limb segment and a distal portion most likely comprising one or more nascent distal tubule segments attached to a GATA3+ collecting duct (Figure 2K).

### Extended organoid culture leads to altered nephron differentiation

Given that d14 organoid nephrons are immature, we were interested to find out whether longer culture times would improve tubule differentiation. It has been suggested that co-labelling with LTL and CDH1 is a sign of proximal tubule maturity (Takasato et al., 2015). Similar to organoid data reported for several other protocols we also observed tubules that were double-positive for LTL and CDH1 in d26 organoids (Morizane and Bonventre, 2017a, Brown et al., 2015). In a subset of d26 organoids, the double-labeled portions were part of large, abnormally branched structures (arrow in Figure 4A). To help interpret these results, we investigated the endogenous expression pattern of LTL and CDH1 in tissue sections from normal human fetal kidneys at week 15-16 of gestation. LTL staining was first apparent in immature proximal tubules at the ‘early capillary loop’ stage and became strongly apical (but not basolateral) in proximal tubules that were identified by co-labeling with Cubilin (CUBN) at the ‘late capillary loop’ stage (Figures 4B, C). Importantly, CDH1 was only found in the distal portions of the developing nephrons. In more mature fetal nephrons, co-labeling of LTL and CDH1 was detected in some tubules (both apically and basolaterally) but only in the descending limb of the loop of Henle in the medulla (Figure S3). We also examined SLC12A3 (distal convoluted tubule) but could not detect it in either ‘early’ or ‘late capillary loop’ stage nephrons, suggesting that this marker appears relatively late in human nephrogenesis.

**Figure 4.**
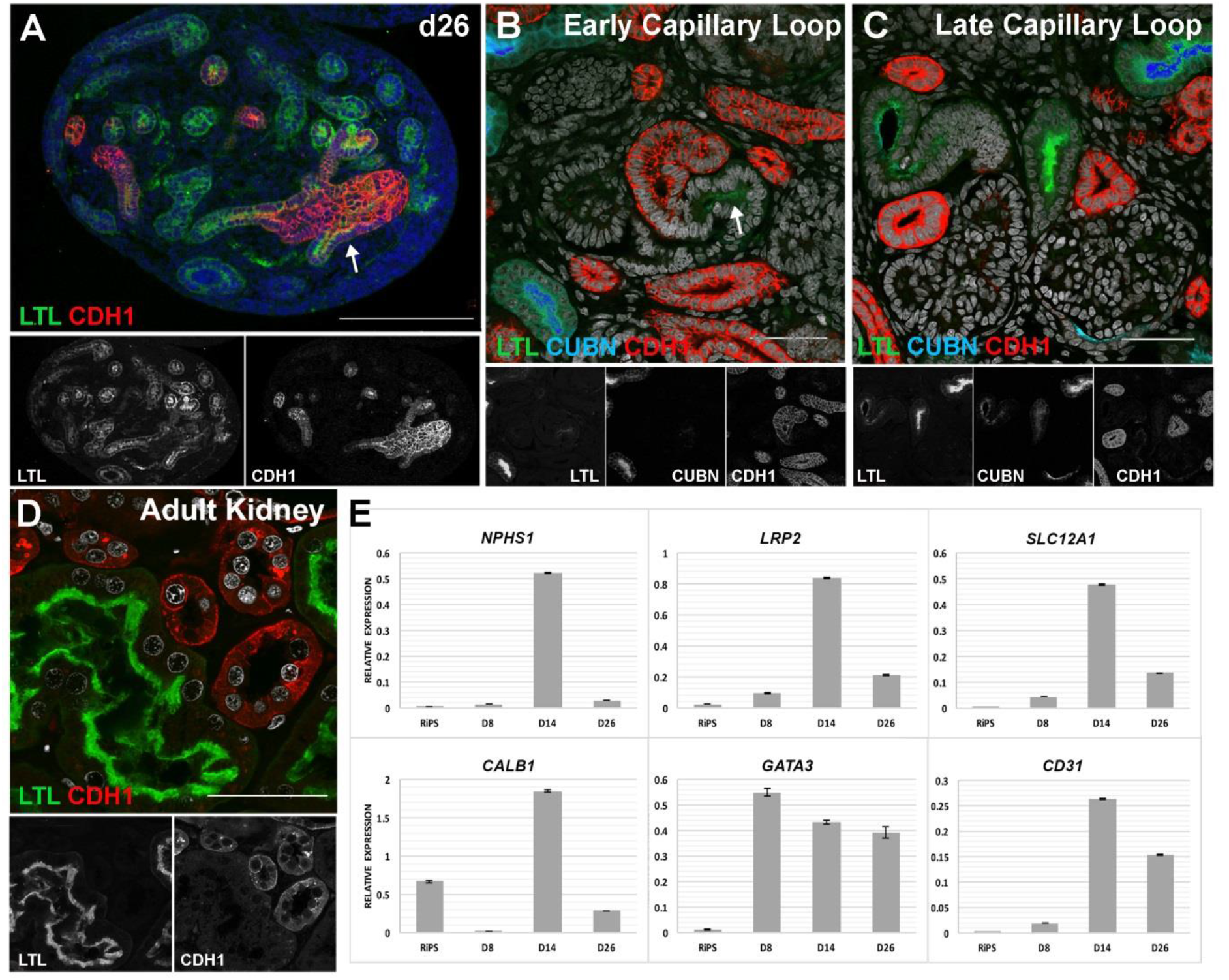
Marker analysis in d26 organoids and human kidney tissue. **(A)** Immunofluorescent staining of a paraffin section of d26 kidney organoid showing partial overlap of LTL and CDH1 in tubules. (**B, C**) Immunostaining of cryosections of human fetal kidney (15-16 weeks gestation) showing LTL^+^ (green; arrow) and CUBULIN^+^ (blue) proximal tubules and CDH1 (red) in distal tubules in ‘early’ (B) and ‘late capillary loop’ (C) stage nephrons (individual color channels are shown below each merged panel). (**D**) Immunofluorescent staining of a paraffin section of an adult human kidney showing non-overlapping staining of LTL (green) and CDH1 (red). (**E**) Quantitative PCR analysis for selected markers in the RiPS line and kidney organoids at d8, d14 and d26. Nuclei are stained with DAPI (blue in (A), white in (B-D)). Scale bars: 50 μm (B-D), 200 μm (A).

To determine if LTL and CDH1 co-label fully mature adult proximal tubules, we analyzed human adult kidney tissue. Similar to the fetal findings, we found that LTL and CDH1 do not overlap, but rather display mutually exclusive labeling of proximal (LTL) and distal (CDH1) portions of the nephrons (Figure 4D). Based on these observations, we conclude that endogenous fetal and adult human proximal tubules do not co-label with LTL and CDH1, thus the mixed (LTL+/CDH1+) identities seen in the organoids may reflect abnormal changes occurring to the tubules following extended culture times. Alternatively this may indicate their conversion to a descending limb-like identity. To further explore whether these changes may be associated with altered differentiation, we measured the expression of the lineage markers *NPHS1, CD31, LRP2, SLC12A1, CALB1* and *GATA3* in whole organoids at d8, d14 and d26 by quantitative polymerase chain reaction (qPCR). While most of these markers increased from d8 to d14, there was a significant downregulation of *NPHS1* (podocytes), *LRP2* (proximal tubule), *SLC12A1* (thick ascending limb) and *CALB1* (distal convoluted tubule, connecting tubule, collecting duct) between d14 and d26 (Figure 4E). A similar downregulation of markers from d14 onwards was observed in the BPEL medium used by Takasato et al. (2015) suggesting that this was not due to our DMEM/KOSR medium (data not shown). The reduction in nephron marker expression at d26 raised the possibility that tubule function was compromised at this stage. To assess this, we performed the fluorescent dextran uptake assay on d26 organoids and found that proximal tubule absorption was significantly impaired (Figure 3G). Taken together, our findings suggest that LTL+/CDH1+ tubules are unlikely to represent mature proximal tubules and that longer organoid culture times beyond d14 result in a downregulation of marker gene expression, loss of proximal tubule absorptive functionality and the formation of potentially abnormal LTL+/CDH1+ tubules. Based on the comparison with human fetal kidneys, our d14 organoid nephrons approximate ‘late capillary loop’ stage nephrons seen in the fetal human kidney (i.e. distinct LTL+ and CDH1+ segments but prior to the onset of *SLC12A3* expression in the distal convoluted tubule segment).

### Organoids at d26 show fibrosis and interstitial cell expansion

When we compared histological sections of d14 and d26 organoids, we noticed that the interstitium around the nephrons was expanded at d26 and was accompanied by accumulation of extracellular matrix and stromal-like cells (data not shown). As this is reminiscent of fibrosis, we performed Masson’s trichrome staining (which stains collagen fibers blue) on d14 and d26 organoid tissue sections. While d14 organoids show little staining, d26 organoids display a fibrotic-like core with collagen staining (Figure 5A). Immunostaining for alpha smooth muscle actin (ASMA), which labels myofibroblasts responsible for the production of fibrotic tissue, revealed a capsule-like layer around d14 organoids but no internal labelling. In contrast, d26 organoids display both capsule-like and interstitial staining, suggesting that between d14 and d26 myofibroblasts arise internally and may be responsible for the deposition of collagen-rich extracellular matrix (Figure 5B). We next compared the MEIS1/2^+^ renal interstitial cells at d14 and d26 and found that this population was qualitatively expanded at d26 (Figure 5C). We noted that between d14 and d26 there was a change in the pattern of proliferation, with d14 organoids showing PCNA^+^ cells in both the nephrons and interstitial cells (Figure 1K) whereas in d26 organoids only rare tubular cells are PCNA^+^ (Figure 5D). Co-labeling of ASMA with MEIS1/2 and PCNA in d26 organoids revealed extensive proliferation of the interstitial cells and a transition to a myofibroblast-like phenotype (Figures 5E, F). In summary, these results suggest that organoid viability is optimal around d14, when the tissue has small amounts of MEIS1/2^+^ interstitial cells and no evidence of fibrosis. After d14, nephron components of the organoid exhibit reduced proliferation while the interstitium continues to expand with some cells converting to an ASMA^+^ myofibroblast-like state that we speculate is responsible for deposition of fibrotic tissue.

**Figure 5.**
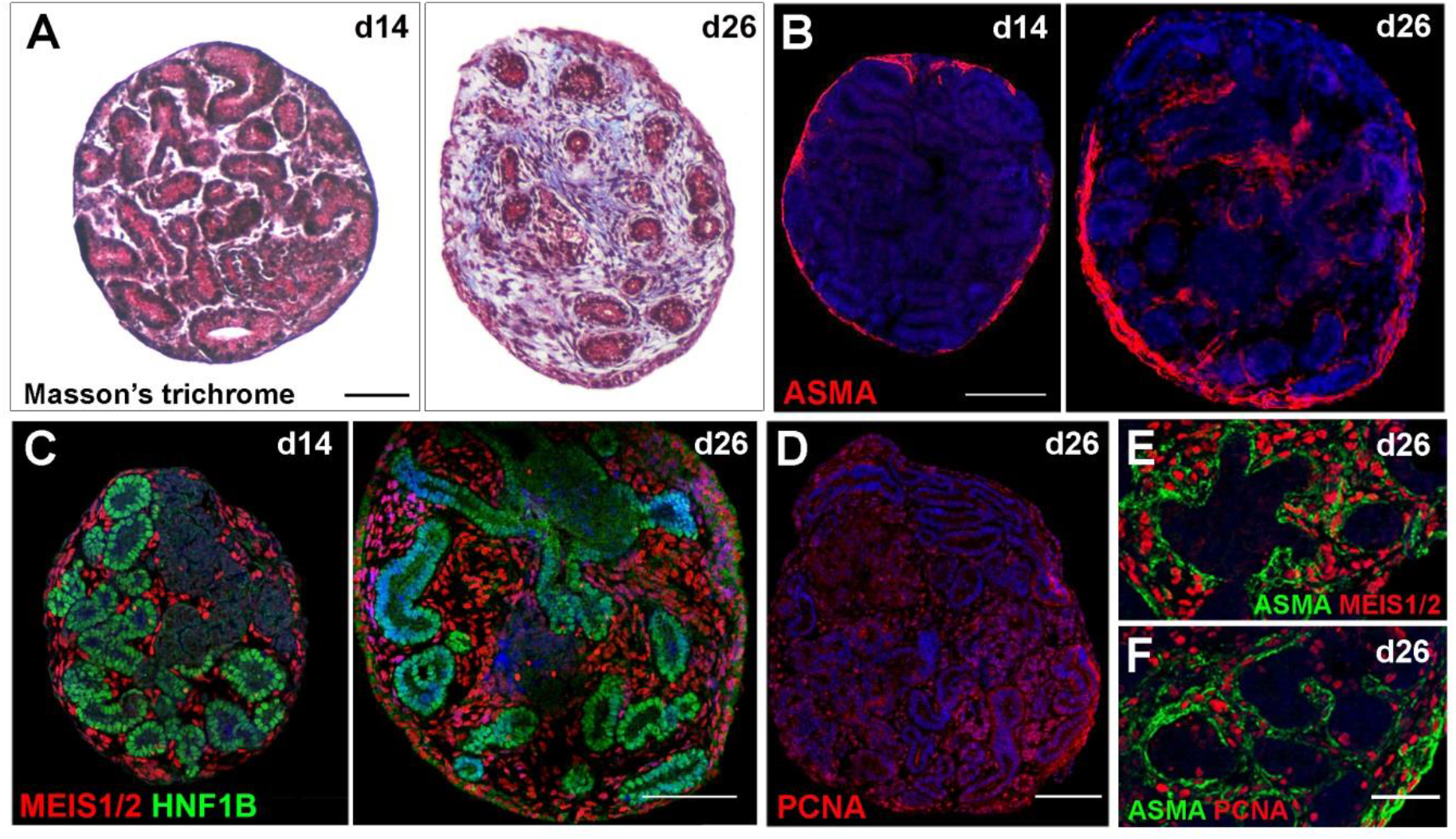
Analysis of fibrosis and interstitial cells in d26 organoids. **(A)** Paraffin sections of d14 and d26 organoids stained with Masson’s trichrome showing collagen deposition at d26 (blue). (**B-F**) Immunofluorescent staining of paraffin sections of d14 and d26 organoids showing **(B)** alpha smooth muscle actin (ASMA)-labeled myofibroblasts (red). **(C)** MEIS1/2^+^ interstitial cells (red) and HNF1B^+^ tubules/ducts (green). **(D)** PCNA^+^ (proliferating) cells (red). (**E, F**) Co-labeling of the myofibroblast marker ASMA (green) with MEIS1/2 (red) in (E) and PCNA (red) in (F) at d26. Nuclei are stained blue with DAPI. Scale bars: 50 μm (E, F), 100 μm (A-D).

### HNF1B knockout organoids as a model of congenital kidney defects

Since the kidney organoids generated with our protocol, as well as those from other methods reported to date, generate fetal human nephrons, we were interested in exploring this system as a way to study human nephrogenesis and genetic renal birth defects. To test this application, we targeted the *HNF1B* transcription factor gene, which has been shown to play essential roles in tubulogenesis in zebrafish and mouse, and when mutated is a common cause of congenital anomalies of the kidney and urinary tract (Nakayama et al., 2010, Naylor et al., 2013, Heliot et al., 2013, Massa et al., 2013, Clissold et al., 2015). Using CRISPR/Cas9 gene editing in MANZ-2-2 iPSCs, we generated two independently-derived iPSC lines with biallelic deletions in exon 2 of the *HNF1B* gene. This mutation, which was confirmed by genomic DNA sequencing, introduces an early translation stop codon and is predicted to result in a non-functional truncated protein that lacks the DNA binding and transactivation domains (Figure 6A). The *HNF1B^−/−^* iPSC lines underwent organoid formation in a similar timeframe as the parental MANZ-2-2 line and with equivalent efficiency (Figure S4). Brightfield imaging of whole organoids showed that by d14, *HNF1B^−/−^* organoids were slightly larger than controls and lacked obvious signs of tubule formation (Figures 6B, C). Expression analysis by qPCR revealed little change in podocyte genes *(NPHS1, WT1)* and *GATA3* but a significant downregulation in markers of the proximal tubule *(LRP2),* thick ascending limb *(UMOD, SLC12A1), CDH1* and *PAX2,* as well as the HNF1B transcriptional targets *HNF4A* and *PKHD2 (UMOD* is also a direct target of HNF1B; Figure 6D). Consistent with these findings, immunostaining of d14 organoids shows a loss of HNF1B protein (a pan-marker of tubules and collecting duct), LTL and SLC12A1 in *HNF1B^−/−^* organoids (Figures 6E-J). Together, these findings indicate that disruption of *HNF1B* leads to a failure in the formation of the proximal tubule and thick ascending limb segments, much like that described in the mouse knockout (Heliot et al., 2013, Massa et al., 2013), with only podocytes and GATA3+ collecting ducts forming in the *HNF1B^−/−^* organoids.

**Figure 6.**
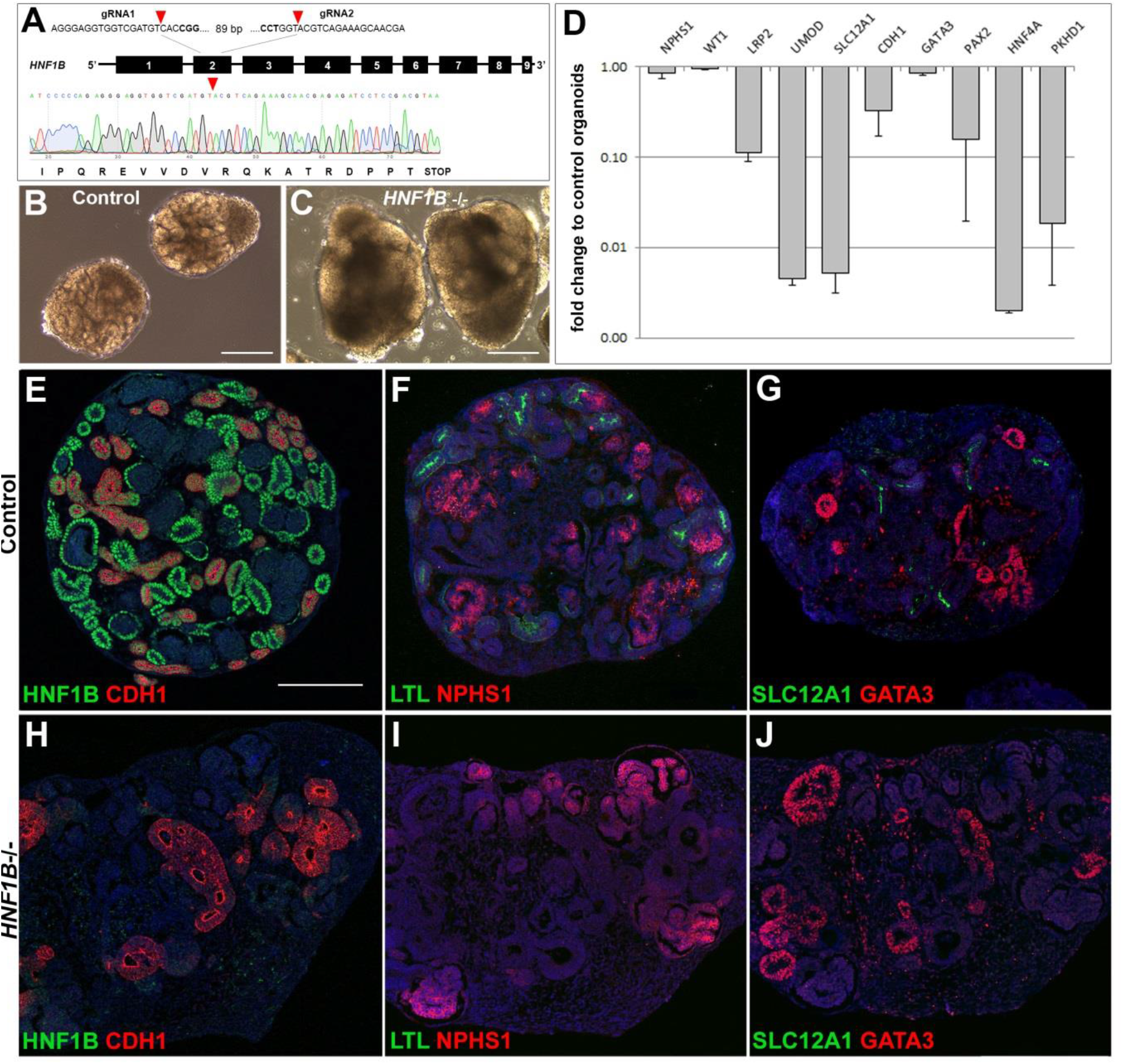
Characterization of HNF1B^−/−^ organoids. **(A)** Schematic overview of the CRISPR/Cas9-based strategy to disrupt the *HNF1B* gene. The extent of the deletion in exon 2 is marked with red arrowheads and the Sanger sequencing chromatogram shows how the resulting frameshift causes a premature stop codon. **(B, C)** Brightfield images of control and *HNF1B^−/−^* organoids at d14. **(D)** Quantitative PCR analysis of selected renal markers and known targets of HNF1B *(HNF4A, PKHD1, UMOD)* in d14 *HNF1B^−/−^* kidney organoids compared to parental control kidney organoids. (E-J) Immunofluorescent staining of paraffin sections of parental control and *HNF1B^−/−^* kidney organoids at d14 for HNF1B (green), LTL (proximal tubule, green), SLC12A1 (thick ascending limb, green), NPHS1 (podocytes, red), CDH1 and GATA3 (distal tubule and collecting duct, red). Nuclei are stained blue with DAPI. Scale bars: 200 μm.

## Discussion

Multiple approaches for differentiating human pluripotent stem cells into kidney organoids have recently been reported (Yamaguchi et al., 2016, Li et al., 2016, Brown et al., 2015, Takasato et al., 2015, Morizane et al., 2015, Freedman et al., 2015, Taguchi et al., 2014). These protocols all result in organoids that successfully recapitulate morphological features of the native fetal kidney but are hampered by cost, technical complexity and reproducibility. These limitations reduce the utility of kidney organoids as a tool for studying basic renal biology, disease/injury modelling and for potential clinical and drug testing applications. By developing a novel EB approach, coupled to an affordable bioreactor culture system that works reliably for diverse pluripotent stem cell lines, we have overcome many of these drawbacks. This simple, scalable and reproducible technology allows the direct formation of large numbers of kidney organoids of defined size and can be readily adopted by non-specialist laboratories and the pharmaceutical industry.

In contrast to most of the existing kidney organoid protocols, which initiate differentiation of pluripotent stem cells in 2D adherent monolayers, we generated 3D aggregates of pluripotent stem cells (aka EBs) that better mimic the structure of the developing embryo (Höpfl et al., 2004). Specifically, the EB environment is thought to be conducive to the formation of endogenous morphogenic gradients and foster homotypic cell sorting and self-assembly of tissues. Such processes are less efficient in 2D monolayer cultures and this may explain the lack of robustness of current methods (Serra et al., 2012). Our protocol gives comparable results with multiple iPSC cell lines that are derived via different reprogramming approaches. Not all of the EBs formed by our method give rise to kidney organoids, with smaller aggregates (<200 um in diameter) often failing to generate tubular tissue. This observation is in keeping with prior work with EBs in which the ‘quality’ of terminal differentiation is dependent on EB size/cell number (Mohr et al., 2010, Messana et al., 2008), presumably because there is a minimal threshold of cells needed to correctly mimic a developing embryo. The range in EB sizes that is generated by our approach is probably the result of differences in the sizes of the iPSC colony fragments, which itself is influenced by the starting confluency and the extent of mechanical dissociation. While further optimization of this step to reduce size heterogeneity is possible, such as fully dissociating the cells and plating them onto uniform microwells (Ungrin et al., 2008, Kim et al., 2014), this has the disadvantage of introducing additional handling steps, cost and places restrictions on large-scale production. Because our approach is easily scalable, it is possible to generate a large number of EBs (we can readily culture ~500 organoids per 150 ml spinner-flask). As a result of this production capacity, size-exclusion via sieving provides a rapid and labor-efficient way of enriching for organoids with optimal renal differentiation potential. This step involves removing specimens with diameters below 200 μm and above 500 μm. This latter group displays signs of core cell death, probably because their size exceeds the limits of oxygen diffusion and mass transport. In support of this, other studies have shown that oxygen diffusion in EBs and organoids is limited to a depth of 200-300 μm, depending on the density and composition of the tissue (Winkle et al., 2012, Hubert et al., 2016). As a result of our size selection step, we can readily obtain EBs with a defined size range that are enriched for kidney tissue, free of core cell death and without the need for laborious handling steps.

A key innovation of our method is the discovery that KOSR can replace more expensive supplements such as B27 used by Freedman et al. (2015) and FGF2/9 used in other protocols (Takasato et al., 2015, Morizane et al., 2015, Li et al., 2016, Taguchi et al., 2014). Preliminary experiments suggest that a minimum of 5 days of KOSR treatment is needed in order to prevent organoid deterioration at later stages (unpublished observations). At this stage it is not clear how KOSR substitutes for the FGF signaling pathway, which acts by maintaining nephron progenitors in an undifferentiated state *in vivo* (Barak et al., 2012). KOSR and B27 both contain transferrin and insulin and these factors are essential for the serum-free culture of fetal kidney explants, where they act synergistically to support growth (Thesleff and Ekblom, 1984, Brewer et al., 1993, Garcia-Gonzalo and Belmonte, 2008, Brière et al., 1991). Whether insulin and transferrin are the key FGF-substituting components in KOSR and B27 remains to be determined. Regardless, our identification of KOSR as an inexpensive ‘Stage II’ medium that supports kidney organoid growth is a major advance since it substantially lowers the cost per assay, thereby enabling kidney organoid generation at an industrial scale. This breakthrough is a critical prerequisite for efforts to develop kidney organoids as a platform for nephrotoxicity testing and for future clinical applications such as cell replacement therapies (Morizane and Bonventre, 2017b).

With regard to technical complexity, our protocol differs from existing methods by being very simple with few handling steps. For instance, the 3D culture method of Morizane et al. (2015), which produces organoids that are morphologically similar to ours, involves additional dissociation and centrifugation steps, as well as laborious growth in 96-well plates. While the most recent Taguchi et al. method requires separate induction of nephron and collecting duct progenitors followed by their co-culture (Taguchi and Nishinakamura, 2017). In our protocol, the EBs are formed in suspension thus enabling them to be size-selected and transferred with ease to spinner-flasks with minimal effort and time. Perhaps as a result of these fewer handling steps, we observe rapid tubule formation with structures visible by brightfield microscopy as early as d8, in contrast to other methods which take between 12-21 days (Chuah and Zink, 2017, Kaminski et al., 2017).

The organoids made from our protocol develop a well-defined proximal-distal segmentation pattern at d14 that is comparable to other reports (Takasato et al., 2015, Morizane et al., 2015, Freedman et al., 2015, Taguchi et al., 2014, Brown et al., 2015, Li et al., 2016). Our direct comparison with fetal human kidney tissue suggests that d14 organoid nephrons are comparable to the ‘late capillary loop’ stage, characterized by an LTL+ proximal segment attached to a CDH1+ distal portion but no morphological signs of the thin limbs of the loop of Henle or expression of *SLC12A3* in a distal convoluted segment. We anticipate that longer culture times might improve the differentiation of the organoid nephrons. While this indeed appears to be the case for podocytes, with the appearance of foot process-like structures at d26, we found that the expression of tubule segment markers decreases after d14 and that overlapping staining of LTL and CDH1 becomes more widespread. Based on observations in the mouse (Cho et al., 1998), it has been suggested that LTL+/CDH1+ tubules represent mature proximal tubules (Takasato et al., 2015). However, in human fetal and adult kidneys we did not observe appreciable overlap with LTL and CDH1, except at a low level in descending limbs in the most mature of the fetal nephrons. While the LTL+/CDH1+ tubules in the organoids may represent differentiation to a descending limb-like identity, given the reduced marker level and the dysmorphic appearance of some of the tubules at d26, we favor the interpretation that it represents an aberrant state. In support of this, human distal tubule cells have a tendency to transdifferentiate into proximal tubule cells in culture (Baer et al., 1999, der Hauwaert et al., 2013), raising the strong possibility that in kidney organoids the CDH1+ distal nephron acquires LTL reactivity (and potentially the expression of other proximal markers). This issue of cell identity is of particular importance to resolve given that nephrotoxins like cisplatin have been found to only induce injury in LTL^+^/CDH1^+^ tubules and not LTL<^+^-only tubules in kidney organoids (Takasato et al., 2015). Similarly, the significance of the common finding that LTL labels both the apical and basolateral domains of organoid proximal tubule cells (Brown et al., 2015, Takasato et al., 2015, Morizane et al., 2015, Freedman et al., 2015), while it is exclusive to the apical domain in human nephrons *in vivo,* needs further investigation.

The reduction in kidney tubule marker gene expression after d14 and the appearance of potentially abnormal LTL+/CDH1+ cells may be related to the fibrotic changes we observed at d26. We found extensive proliferation of MEIS1/2^+^ interstitial cells with some co-labeling with ASMA, a hallmark of pro-fibrotic myofibroblasts, and the deposition of collagen-rich extracellular matrix. Renal fibrosis, defined as the disproportionate accumulation of extracellular matrix and the proliferation of interstitial fibroblasts, is one of the important pathways involved in end-stage renal failure and is thought to result from a normal wound-healing response becoming deregulated (Humphreys, 2017). It is possible that extended culture beyond d14 leads to tubular injury, perhaps in response to limitations in the mass transport of oxygen or nutrients into the organoid. While we sieve out large organoids with overt core apoptosis, the smaller organoids may still be undergoing sub-lethal levels of stress and releasing pro-fibrotic signals to the interstitium, via tubulointerstitial crosstalk pathways (Maarouf et al., 2016). Proliferative changes in the growth of the tubules and interstitium also coincides with the fibrotic phenotype, with both populations dividing at d14 whereas at d26, growth is largely restricted to MEIS1/2^+^/ASMA^+^ cells. Large numbers of stromal cells are seen in kidney organoids reported by other methods, suggesting that this issue of interstitial expansion may be a widespread phenomenon (Takasato et al., 2015, Morizane and Bonventre, 2017a). As the pathogenesis of renal fibrosis remains poorly understood, and is complicated by the involvement of the immune system in animal models, kidney organoids may provide a useful tool to dissect the non-immune signals involved in fibrogenesis and the screening of anti-fibrotic drugs.

We conclude that d14 organoids derived from our protocol provide an optimal timepoint to study human nephrogenesis with the generation of tissue that resembles ‘late capillary stage’ nephrons and is free of fibrosis. To demonstrate the utility of our protocol to study human kidney development, we used gene editing in iPSCs to disrupt the *HNF1B* gene, which is implicated in congenital anomalies of the kidney and urinary tract in 10% of cases in humans (Nakayama et al., 2010). Kidney organoids made from *HNF1B^−/−^* iPSCs contained podocytes and GATA3^+^ presumptive collecting ducts but strikingly lacked cells with either proximal or distal nephron identities. This phenotype closely resembles the rudimentary nephrons observed in *HNF1B* conditionally-deficient mice, where glomeruli are found connected to the collecting duct system via a short primitive tubule (Heliot et al., 2013, Massa et al., 2013). Our phenocopy of this result validates the use of kidney organoids as a human-based system to study renal gene function. *hnf1b^−/−^* mice (and individuals with heterozygous mutations in *HNF1B*) also develop renal cysts but these were not observed in our *HNF1B^−/−^* kidney organoids at d14. Cystic phenotypes likely require longer culture time to develop. Indeed, in a similar study, where kidney organoids were used to recapitulate polycystic kidney disease, cyst formation was only apparent after 5 weeks in culture (Cruz et al., 2017, Freedman et al., 2015).

Looking forward, further improvements to existing kidney organoid protocols are needed if the resulting nephrons are to stably maintain their identity and mature beyond fetal stages. While kidney organoids have value for understanding kidney developmental biology, more mature tissue will be better suited to model adult renal diseases and acute injury. It is generally thought that improved vascularization of the organoids will aid in nephron maturation by providing better perfusion of the tissue and developmental signals, such as those coming from peritubular capillary cells and fluid flow down the nephron. Strategies to overcome these issues include the transplantation of organoids into permissible sites of immune-deficient mice where recruitment and vascularization by host blood vessels can occur (Sharmin et al., 2016, Francipane and Lagasse, 2016) and the culture of organoids in microfluidic devices (‘organs-on-chips’; Bhatia and Ingber, 2014, Wilmer et al., 2016). However, testing of such strategies will require a large and steady supply of uniform kidney organoids and this is a clear bottleneck with current protocols. The EB-based approach for generating kidney organoids described here is simple, rapid, scalable, and works robustly for a range of different pluripotent stem cell lines. Our method overcomes many of the drawbacks hampering existing protocols and provides the advance needed in order for kidney organoids to be effectively used for future applications such as drug testing, optimization of vascularization, and cell replacement therapies where there is a requirement to generate large amounts of tissue.

## Methods

All experiments were performed with the three iPSC lines described below. For immunohistochemistry, electron microscopy and qPCR, data are representative of all three lines (unless stated otherwise). Individual stainings and measurements were performed at least in triplicate.

### iPSC lines and maintenance

All work was carried out with the approval of the University of Auckland Human Participants Ethics Committee (UAHPEC 8712) and biosafety approval (GMO05). BJ RiPS (reprogrammed by RNA) were a gift from Dr Chad Cowan (Warren et al., 2010). CRL1502 clone C32 (reprogrammed by episomal vectors) were developed in the Wolvetang laboratory (Briggs et al., 2013). The MANZ-2-2 line was generated in the Davidson laboratory. Briefly, the MANZ2-2 line was generated from participant skin biopsies that were dissected and treated with 0.5% trypsin/EDTA solution. Cells were rinsed with DPBS and seeded into fibroblast growth medium (DMEM/15% FBS, 1% penicillin/streptomycin) on gelatin coated plates. Resulting outgrowths of dermal fibroblasts were expanded and reprogrammed using the CytoTune™-iPS 2.0 Sendai Reprogramming Kit (Invitrogen) according to the manufacturer’s instructions. On day 7, the cells were dissociated with 0.05% trypsin, washed and seeded into fibroblast growth medium on LDEV-free hESC-qualified Geltrex-coated dishes. On day 8 the medium was changed to TeSR-E8 (STEMCELL Technologies). Cultures were monitored for emerging iPSC colonies for 3-4 weeks post transduction. Clonal colonies were manually picked and expanded as individual iPSC lines. New iPSC lines were karyotyped by LabPLUS (Auckland District Health Board) and analyzed for expression of pluripotency markers by qPCR and immunocytochemistry. Established iPSC lines were maintained on Geltrex-coated dishes in mTeSR1 (STEMCELL Technologies) plus 1% penicillin/streptomycin and 2.5 μg/mL Plasmocin (InvivoGen). At ~70% confluence, cells were dissociated using 1/3 Accutase in DPBS. Cells were scraped from the dish, pelleted at 800 rpm for 5 min and resuspended in mTeSR1 plus 5 μM Y27632 for the first 24 hours to facilitate cell survival.

### Organoid formation

Prior to EB formation, iPSCs were cultured on a 10 cm Geltrex-coated dish to 40-50% confluency when discrete colonies have formed. Cells were washed twice with DPBS and incubated with 2 ml of 1 mg/ml Dispase solution for 6 min at 37°C. Cells were washed three times with DBPS, scraped with a cell lifter, resuspended in BPEL (Ng et al., 2008) plus 8 μM CHIR99021, 3 μM Y27632 and 1 mM beta-mercaptoethanol and evenly distributed into ultra-low 6-well attachment plates (Corning). Half medium change was carried out on day 2 with BPEL supplemented with 8 μM CHIR99021. On day 3, EBs were allowed to sediment in a 50 mL tube and washed twice in DMEM. EBs were returned to the ultra-low 6-well attachment plate and transferred to Stage II culture medium (DMEM, 15% KnockOut Serum Replacement (KOSR; Thermo Fisher), 1% NEAA, 1% penicillin/streptomycin, 1% HEPES, 1% GlutaMAX, 0.05% PVA, 2.5 μg/mL Plasmocin). Once tubule formation was observed (day 7-8), organoids were transferred into a 150 mL spinner flask (Corning) in 45 mL of Stage II medium, stirring at 90 rpm, with half medium change every 2 days. Alternatively, organoids were left in the ultra-low 6-well attachment plate, and agitated using an orbital shaker (Hubert et al., 2016). No differences in organoid size and shape were observed when grown in either spinner flasks or plates.

### Size filtration and measurement

Organoids were passed through 200 and 500 μm strainers (pluriStrainer). Whole organoid photos were acquired on an EVOS XL inverted microscope. Efficiency of tubule formation and size measurements were performed using ImageJ.

### Reabsorption assay

20 μg/ml 10kDa Rhodamine-dextran was added to Stage II culture medium for 48 hrs. Organoids were washed in Stage II medium for 5 hrs before fixation in 4% PFA, paraffin embedding and sectioning.

### HNF1B knock-out

gRNA pairs targeted to introduce an 89 bp deletion in exon 2 of the *HNF1B* gene were designed using the RGEN, COSMID and CCtop online tools (http://www.rgenome.net/cas-designer/, (Park et al., 2015); https://crispr.bme.gatech.edu/, (Cradick et al., 2014); https://crispr.cos.uni-heidelberg.de/, (Stemmer et al., 2015). gRNAs were cloned into the pSpCas9(BB)-2A-GFP (Addgene 48138) construct and knockout efficiencies for gRNA pairs were evaluated in HEK293 cells. Plasmids containing the gRNAs with highest knockout efficiency (HNF1B_gRNA1: 5’-AGGGAGGTGGTCGATGTCACCGG-3’; HNF1B_gRNA2: 5’-CCTGGTACGTCAGAAAGCAACGA-3’) were introduced into the MANZ-2-2 iPSC line by reverse transfection using TransIT-LT1 (Mirus Bio). 48 hrs post transfection, GFP-positive cells were isolated by flow cytometric sorting and 8,000 cells were plated on a 10 cm Geltrex-coated dish into pre-warmed mTeSR1 plus 5 μM Y27632. Medium changes were carried out daily using conditioned mTeSR1 without Y27632. Single colonies were manually picked when they had reached a suitable size (~10 days post plating), clonally expanded, and screened for biallelic deletion clones using PCR primers flanking the deleted region (Peters et al., 2013). Homozygote deletions were verified by Sanger sequencing. Clones were expanded, karyotyped and used for organoid assays. Control experiments were carried out with MANZ-2-2 iPSCs.

### RNA extraction, cDNA synthesis and qPCR

Organoids were briefly washed in DPBS before transfer into TRIzol. RNA was extracted using the GENEzol TriRNA Pure kit (Geneaid). cDNA was synthesized using qScript cDNA SuperMix (Quanta). For qPCR, PerfeCTa SYBR Green FastMix (Quanta) was used. qPCR was performed on a QuantStudio 6 Flex Real-Time PCR machine. Primers used are listed in Table S1. Samples were normalized to *HPRT1* expression. Gene expression was calculated using the ddCt method. Error bars represent standard deviation from triplicates.

### Histology and Immunochemistry

Organoids were fixed in 4% paraformaldehyde/PBS (PFA) overnight at 4°C. After washing with PBS plus 0.1% Tween, organoids were transferred into an embedding mold and filled with embedding medium (1% low-melting agarose, 0.9% agar, 5% sucrose). Once solidified, the blocks were transferred into 70% ethanol and incubated 4°C overnight. Over the next two days, the blocks were transferred through a series of 95% and 2x 100% ethanol, 50:50 ethanol:xylol, 100% xylol, 1 hr each, rocking at room temperature, followed by 50:50 xylol:paraffin at 65°C overnight and changes of paraffin every 4 hrs. After embedding, the blocks were sectioned at 6 μm on a Leica microtome. Sections were air-dried, then stored at 4°C. Human week 15-16 fetal kidneys were fixed in 4% PFA overnight at 4°C. After washing twice with PBS the samples were incubated in 30% sucrose in PBS at 4°C for 36 hrs (till the tissue had sunk to the bottom of the vials), then embedded in Optimal Cutting Temperature (OCT, Tissue-Tek) cryoblocks. The cryo-preserved samples were sectioned at 10 μm and stored at −80°C. For H&E staining, paraffin sections were deparaffinized at 65°C for 30 min, then incubated in two changes of xylol (10 min each). H&E staining was performed following standard procedures and imaged on a Leica DMIL LED microscope. Immunohistochemistry was performed using standard procedures. Antigen retrieval (10 mM sodium citrate buffer plus 0.05% Tween-20, pH 6.0 at 95°C for 30 min) was carried out for all antibodies except CASP3. Immunostainings were imaged using Zeiss a LSM710 or Leica SP8 confocal microscopes. List of antibodies see Table S2.

### Electron microscopy

Organoids were fixed in 2.5% glutaraldehyde, 2% PFA and 0.1 M phosphate buffer pH 7.4 at 4°C, and kept in the fixative until processing. Samples were washed in 0.1 M Sörensens phosphate buffer 3x 10 min, post-fixed in 1% osmium tetroxide in 0.1 M phosphate buffer for 1 hr at room temperature and washed 2x in 0.1 M phosphate buffer for 5 min. The samples were dehydrated in a series of ethanol washes, 10 min each at room temperature (30%, 50%, 70%, 90% and 2x 100%), followed by 2x propylene oxide wash for 10 min at room temperature. All washes were performed on an orbital shaker. The samples were infiltrated with a graded series of propylene oxide:epoxy 812 resin mix (1:1) for 1 hour, before embedding in freshly made pure resin overnight. The samples were then placed into molds and polymerized at 60°C for 48 hrs. Ultra-thin sections of 70 nm were placed on copper 200 mesh grids and stained with uranyl acetate and lead citrate. Sections were imaged using Tecna G^2^ Spirit Twin transmission electron microscope.

### Fetal human tissue

Consented, de-identified, human fetal tissue from elective terminations was collected following review of the study by the Institutional Review Board at Keck School of Medicine of the University of Southern California. Week 15-16 kidney samples were supplied by collaborators at the Children’s Hospital of Los Angeles, the University of California, San Francisco or the Wellcome-funded Human Developmental Biology Resource center at the Institute of Genetic Medicine, Newcastle Upon Tyne, UK and the Institute of Child Health, London, UK. Gestational age identification followed the guidelines of the American College of Obstetricians and Gynecologists. Samples from the Children’s Hospital of Los Angeles were transported on ice at 4°C in 10% fetal bovine serum, 25 mM HEPES, high glucose DMEM and delivered within 1-3 hrs after procedures. Samples from the University of California, San Francisco were transported in similar media and delivered on the next day after the procedure. Samples from the Human Developmental Biology Resource center were supplied as whole tissues fixed in 4% PFA and shipped in PBS, or as paraffin sections or whole tissue embedded in paraffin.

## Acknowledgements

We thank Adrian Turner, Jacqui Ross and Satya Amirapu for help with histology and microscopy, Paula Lewis for critical reading of the manuscript, and Peter Shepherd for providing resources and personnel to facilitate the project. Work in A. J. D.’s laboratory was supported by Health Research Council New Zealand (17/425), Auckland Medical Research Foundation (1116018), Cystinosis Research Foundation USA, Cystinosis Research Ireland Foundation, Maurice Wilkins Centre, Valrae Collins philanthropic support for A. P. Work in A. P. M.’s laboratory was supported by the National Institutes of Health (DK054364) and California Institute of Regenerative Medicine (LA1-06536).

## Author contribution

A.P., T.M.H and A.J.D. conceptualized the study, A.P. and VS. designed the experiments, A.P. established the protocol, A.P. and VS. conducted organoid culture, processing and analysis, VS. generated the *HNF1B* knockout, T.T. performed immunohistochemistry on fetal kidney tissue, A.P., J.A.H., J-H.S., B.S. and T.M.H. generated the MANZ-2-2 iPSC line, E.J.W. generated the C32 iPSC line, A.P.M. and T.M.H co-supervised the study, A.P., V.S. and A.J.D. wrote the manuscript, A.J.D supervised the study, and VS. and A.J.D acquired funding.

